# Orphan nuclear receptor COUP-TFII drives the myofibroblast metabolic shift leading to fibrosis

**DOI:** 10.1101/2020.06.05.135889

**Authors:** Li Li, Pierre Galichon, Xiaoyan Xiao, Ana C Figueroa-Ramirez, Diana Tamayo, Jake June-Koo Lee, Marian Kalocsay, David Gonzalez-Sanchez, Maria S Chancay, Kyle McCracken, Dario Lemos, Nathan Lee, Takaharu Ichimura, Yutaro Mori, M. Todd Valerius, Xiaoming Sun, Elazer R Edelman, Joseph V Bonventre

## Abstract

Recent studies demonstrated that metabolic disturbance, such as augmented glycolysis, contributes to fibrosis. The molecular regulation of this metabolic perturbation in fibrosis, however, has been elusive. COUP-TFII (also known as NR2F2) is an important regulator of glucose and lipid metabolism. Its contribution to organ fibrosis is undefined. Here, we found increased COUP-TFII expression in myofibroblasts in kidneys of patients with chronic kidney disease, fibrotic lungs of patients with idiopathic pulmonary fibrosis, fibrotic human kidney organoids, and fibrotic mouse kidneys after injury. Genetic ablation of COUP-TFII in mice resulted in attenuation of injury-induced kidney fibrosis. A non-biased proteomic study revealed the suppression of fatty acid oxidation and the enhancement of glycolysis pathways in COUP-TFII overexpressing fibroblasts. Overexpression of COUP-TFII in fibroblasts was sufficient to enhance glycolysis and increase alpha smooth muscle actin (αSMA) and collagen1 levels. Knockout of COUP-TFII decreased glycolysis and collagen1 levels in fibroblasts. Chip-qPCR assays revealed the binding of COUP-TFII on the promoter of PGC1α, a critical regulator of mitochondrial genesis and oxidative metabolism. Overexpression of COUP-TFII reduced the cellular level of PGC1α. In conclusion, COUP-TFII mediates fibrosis by serving as a key regulator of the shift in cellular metabolism of interstitial pericytes/fibroblasts from oxidative respiration to aerobic glycolysis. The fibrogenic response may share a common pathway in different organ injury and failure. Targeting COUP-TFII serves as a novel treatment approach for mitigating fibrosis in chronic kidney disease and potential other organ fibrosis.

## Introduction

Kidney fibrosis is a pathologic hallmark of CKD, affecting approximately 10% of the world’s adult population (Jha, Garcia-Garcia et al., 2013, Romagnani, Remuzzi et al., 2017). The progressive deposition and expansion of the fibrotic matrix in kidney parenchyma ultimately lead to kidney failure (Bonventre & Yang, 2011, Rockey, Bell et al., 2015). A similar process was found in other fibrotic diseases, such as liver cirrhosis, idiopathic pulmonary fibrosis, and scleroderma (Wynn & Ramalingam, 2012). The global burden of fibrosis is high, with an estimated prevalence rate at 1 in 4 people (Zhao, Kwan et al., 2019). Currently, there is no cure for fibrosis, highlighting the need for a new strategy and a better understanding of the molecular mechanisms underlying fibrosis. It is generally accepted that myofibroblasts are major cellular contributors to fibrotic disease (Bonventre & Yang, 2011, Duffield, 2014, Duffield, Lupher et al., 2013, Tomasek, Gabbiani et al., 2002). Myofibroblasts express alpha smooth muscle actin (αSMA) and contract, migrate and produce excessive extracellular matrix (ECM) (Klingberg, Hinz et al., 2013). Comprehensive genetic fate mapping studies in rodents indicate that kidney resident *Foxd1* expressing pericytes/perivascular fibroblasts/mesenchymal stem cell-like stromal cells are the main source of myofibroblasts after kidney injury (Falke, Gholizadeh et al., 2015, Humphreys, Lin et al., 2010, Humphreys, Valerius et al., 2008, Kobayashi, Mugford et al., 2014, Kramann, Schneider et al., 2015, Lin, Kisseleva et al., 2008, Picard, Baum et al., 2008).

Recently, an emerging body of evidence has demonstrated the link between metabolic dysregulation and fibrosis (Hou & Syn, 2018, Lan, Geng et al., 2016, Xie, Tan et al., 2015, Zank, Bueno et al., 2018). Genome-wide transcriptome profiling revealed inflammation and metabolism as the top dysregulated pathways in fibrotic human kidneys (Kang, Ahn et al., 2015). A similar analysis of human skin fibrosis identified perturbations of fatty acid oxidation (FAO) and glycolysis pathways (Zhao et al., 2019). Inhibition of glycolysis or restoring FAO by genetic or pharmacological methods demonstrated promising results to mitigate fibrosis in various animal models (Ding, Jiang et al., 2017, Han, Wu et al., 2017, Kang et al., 2015, Tran, Zsengeller et al., 2016, Zhao et al., 2019). Despite these preliminary findings, the exact mechanisms that regulate metabolic dysregulation, especially in myofibroblast, remain largely unknown.

Chicken ovalbumin upstream promoter-transcription Factor II (COUP-TFII, also known as *NR2F2*) is an orphan member of the nuclear receptor family with unknown endogenous ligands (Pereira, Qiu et al., 1999). Since COUP-TFII has been reported to regulate metabolic functions (Ashraf, Sanchez et al., 2019, Li, Xie et al., 2009, Planchais, Boutant et al., 2015), we evaluated whether it played an important role in fibrosis. Downstream proteins under COUP-TFII control tend to be involved in energy production, anabolic pathways, and cell cycle progression, all of which impact on cell proliferation (Chen, Qin et al., 2012, Planchais et al., 2015, Wu, Kao et al., 2015). Upregulation of COUP-TFII in numerous cancers, such as colon (Bao, Gu et al., 2014), pancreas (Polvani, Tarocchi et al., 2014), prostate (Qin, Wu et al., 2013), and renal cell carcinoma (Fang, Liu et al., 2020), further support the role of COUP-TFII in promoting cell proliferation. We hypothesized that it contributed to organ fibrosis via a regulatory role in the metabolism of the myofibroblast.

In this study, we demonstrate that COUP-TFII is a key regulator of shifting myofibroblast metabolism towards enhanced glycolysis with generation of pro-fibrotic mediators. In our studies increased COUP-TFII expression co-localized with αSMA expression in stromal cells of fibrotic human kidneys, lungs, kidney organoids, and fibrotic mouse kidneys after injury. Ablation of COUP-TFII in adult mice attenuated injury-induced kidney fibrosis. Our findings demonstrate a previously unrecognized role of COUP-TFII on regulating myofibroblast differentiation and highlight it as a relevant therapeutic target to prevent organ fibrosis after injury.

## Results

### COUP-TFII expression is increased in myofibroblasts in human fibrotic diseases, and human kidney organoids

We interrogated previously published, human CKD microarray datasets and found a significant increase in COUP-TFII mRNA levels (1.9-fold) in renal biopsy tissues of 53 patients with CKD (GSE66494) (Fig. 1c) (Nakagawa, Nishihara et al., 2015). In another large cohort (n=95) of microdissected human kidney samples from diabetic or hypertensive CKD subjects with pathology-defined fibrosis, microarray transcription profiling also revealed a significant up-regulated COUP-TFII by analysis of total kidney mRNA levels (data not shown) (Kang et al., 2015). Results from these two independent cohorts of patients suggest an association between COUP-TFII mRNA expression and CKD in humans. We confirmed COUP-TFII protein expression and localization by immunofluorescent staining of normal and diseased human kidneys. In control ‘healthy’ kidney, which was obtained from the non-tumor portions of total nephrectomy samples in patients with renal cell carcinoma, we found little scattered COUP-TFII expression (Fig 1a). In the setting of kidney injury (patients with either acute thrombotic microangiopathy (TMA) or chronic diabetic nephropathy (DN)), however, the number of COUP-TFII-positive cells was significantly increased (Fig 1a). The majority of these cells were localized in the interstitial region and co-localized with expanded αSMA-positive areas of fibrosis (Fig. 1a). Similar results were also identified in human fibrotic lung from patient with idiopathic pulmonary fibrosis (IPF) (Fig 1b).

**Figure 1.**
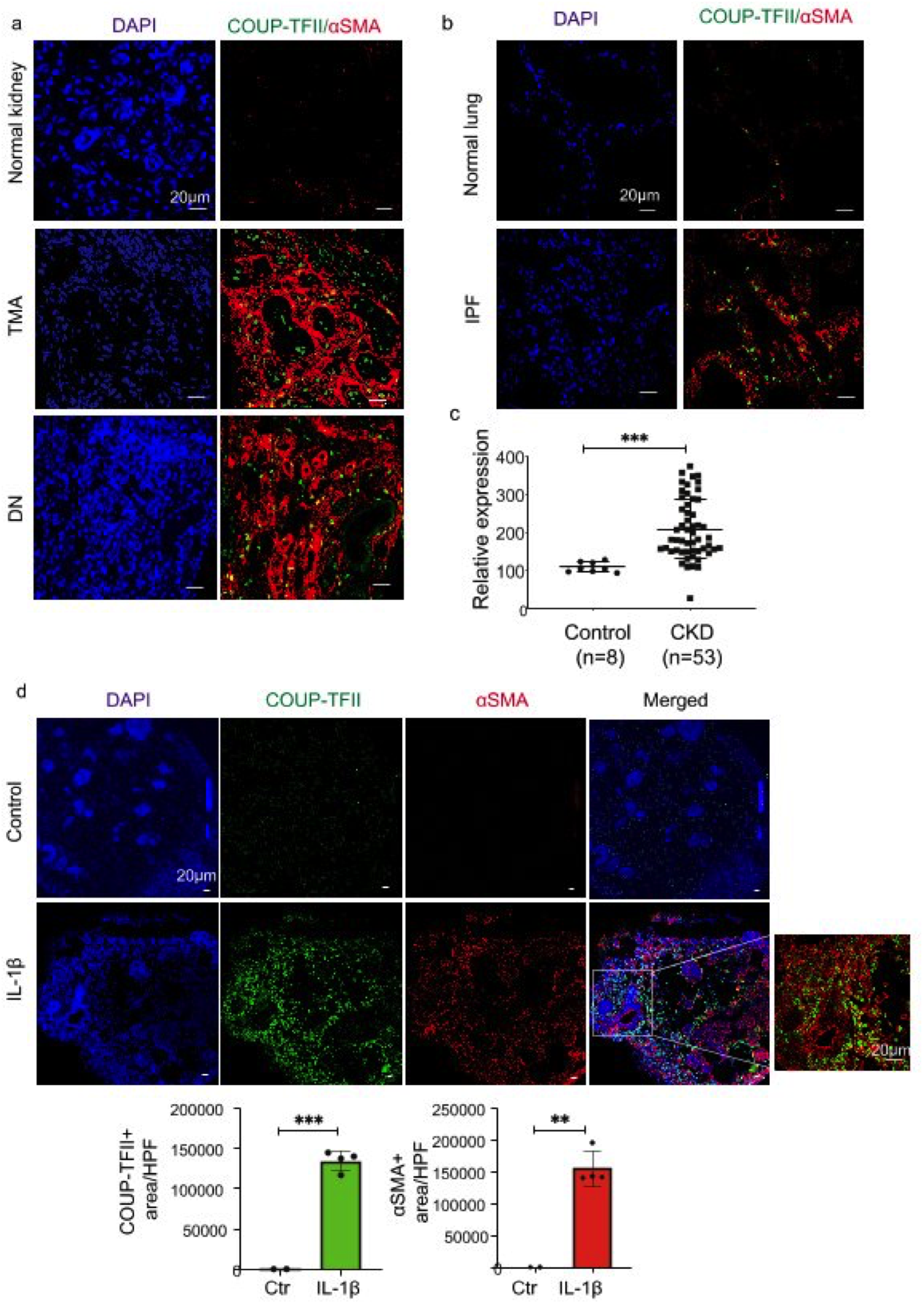
COUP-TFII expression is increased in human fibrotic kidneys and lungs, and IL-1β treated human kidney organoids. COUP-TFII expression is increased in (a) human kidneys from patients with TMA and diabetic nephropathy and (b) in lungs from patients with IPF by immunofluorescence. (c) COUP-TFII mRNA levels in human kidney tissues of control (n=8) and CKD (n=53) subjects from GSE66494. ***p<0.001 by t-test. (d) Human kidney organoids were treated with IL-1β (10ng/ml) for 96 hours. Immunofluorescence reveals significantly increased COUP-TFII and αSMA expression in IL-1β treated organoids, compared to nontreated organoids. Quantification by confocal micrographs in 200x hpf. ****p<0.0001 by t test; mean ± SD.

Next, we examined COUP-TFII expression in human kidney organoids generated by directed differentiation of human pluripotent stem cells (Morizane, Lam et al., 2015). Human kidney organoids provide advantages of 3D nephron structures with multiple human kidney cell types and a rich stroma. Fibrosis of human kidney organoids was induced by incubation with IL-1β for 96 hours as previous reported (Lemos, McMurdo et al., 2018). We then performed immunostaining of COUP-TFII on control and IL-1β treated organoids. As showed in Fig 1d, COUP-TFII expression significantly increased in IL-1β treated organoids compared to control. Most COUP-TFII positive cells were located in the interstitial region and co-localized with αSMA, as we observed in the kidneys of human subjects with CKD. Together, these results suggest an association of increased COUP-TFII expression with fibrosis in humans. The potential link to fibrosis prompted us to investigate the spatial and temporal expression, and mechanistic implications of COUP-TFII in non-injured and injured kidneys in adult mice.

### COUP-TFII protein is expressed in stromal cells in non-injured and injured mouse kidneys

In healthy adult mouse kidneys, a few scattered COUP-TFII positive cells were present within the interstitium. The majority of COUP-TFII positive cells expressed PDGFR-β, a pericyte/fibroblast marker. COUP-TFII was not observed in endothelial cells marked by CD31 (Fig 2a).

**Figure 2.**
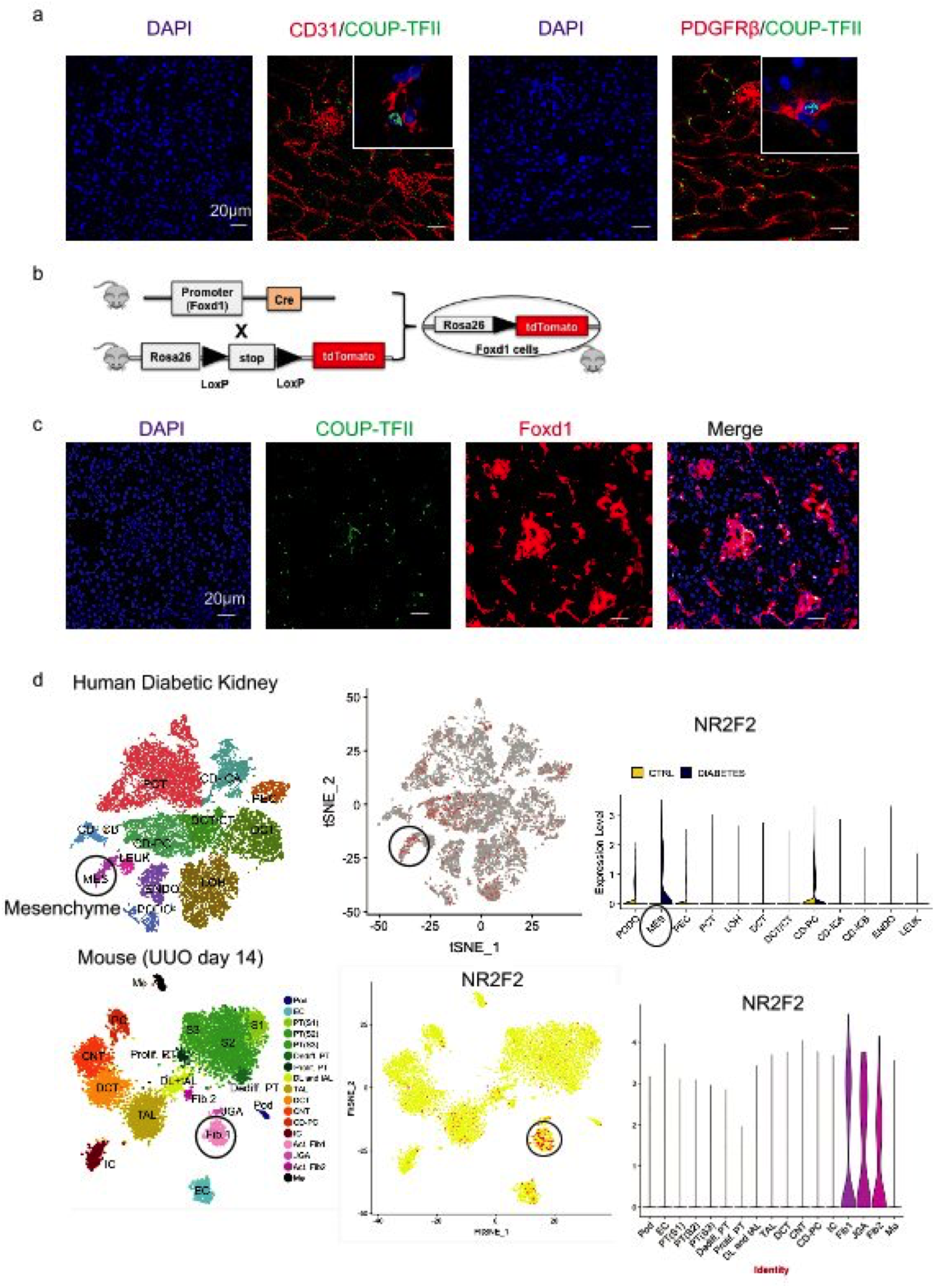
COUP-TFII protein is expressed in pericytes/fibroblasts, both in non-injured and injured mouse kidneys. (a) COUP-TFII+ cells (green) express pericyte/fibroblast marker (PDGFRβ) but not the endothelial cell marker (CD31). (b) Strategy of genetically labeling *Foxd1*-derived stromal cells in non-injured mouse kidney. (c) Most COUP-TFII expression cells (green) overlap with tdTomato-labeled *Foxd1*-derived stromal cells (red) in the non-injured mice kidney. (d) COUP-TFII was enriched most in pericytes/fibroblasts in injured kidney, both in the human (Diabetic kidneys, Fig 2d top) and mouse model (UUO day 14, Fig 2d bottom) as analyzed from single cell sequencing database (http://humphreyslab.com/SingleCell/).

To further assess the developmental origin of COUP-TFII+ cells, we crossed *Foxd1-Cre* driver mice (Humphreys et al., 2010) to tdTomato reporter mice (Madisen, Zwingman et al., 2010) to genetically label the *Foxd1*-derived stromal cells (Fig 2b). The majority of COUP-TFII protein expressing cells (green) overlapped with tdTomato-labeled *Foxd1*-derived stromal cells (red) (Fig 2c), demonstrating that COUP-TFII+ cells in non-injured mice derive from the *Foxd1* population. Analysis of an available single-cell RNA sequencing database (http://humphreyslab.com/SingleCell/) confirmed that COUP-TFII RNA was most enriched in pericyte/fibroblast cells in injured kidneys, both in human (diabetic kidney) (Fig 2d top) and mouse (UUO day 14) (Fig 2d bottom). Thus the majority of COUP-TFII+ cells are kidney stromal cells, both in non-injured and injured kidneys.

### COUP-TFII expression is increased during the development of kidney fibrosis in various kidney injury models in mice

COUP-TFII expression was significantly increased and co-localized in αSMA+ cells within fibrotic regions in the injured kidney in two mouse kidney injury models: unilateral ureteral obstruction (UUO) and unilateral ischemia reperfusion injury (UIRI) (Fig 3a). To further characterize the spatial and temporal expression kinetics of COUP-TFII, we focused on the UUO model since it more reliably induces fibrosis in a short time frame. As shown in Fig 3b, COUP-TFII expression was upregulated as early as day 2 after injury, which is well before upregulation of αSMA and any histologic evidence of fibrosis was evident. We further demonstrated that the increased COUP-TFII expression is distinct from the inflammatory infiltrate that accompanies injury and fibrosis, as it does not overlap with markers of T lymphocytes (CD3), neutrophils (LyG6), or macrophages (F4/80) (Fig 3c). Thus we conclude that COUP-TFII is upregulated specifically within the stromal compartment following kidney injury, and it precedes the expression of fibrotic markers, indicating a potential causative role in the pathophysiology of fibrosis.

**Figure 3.**
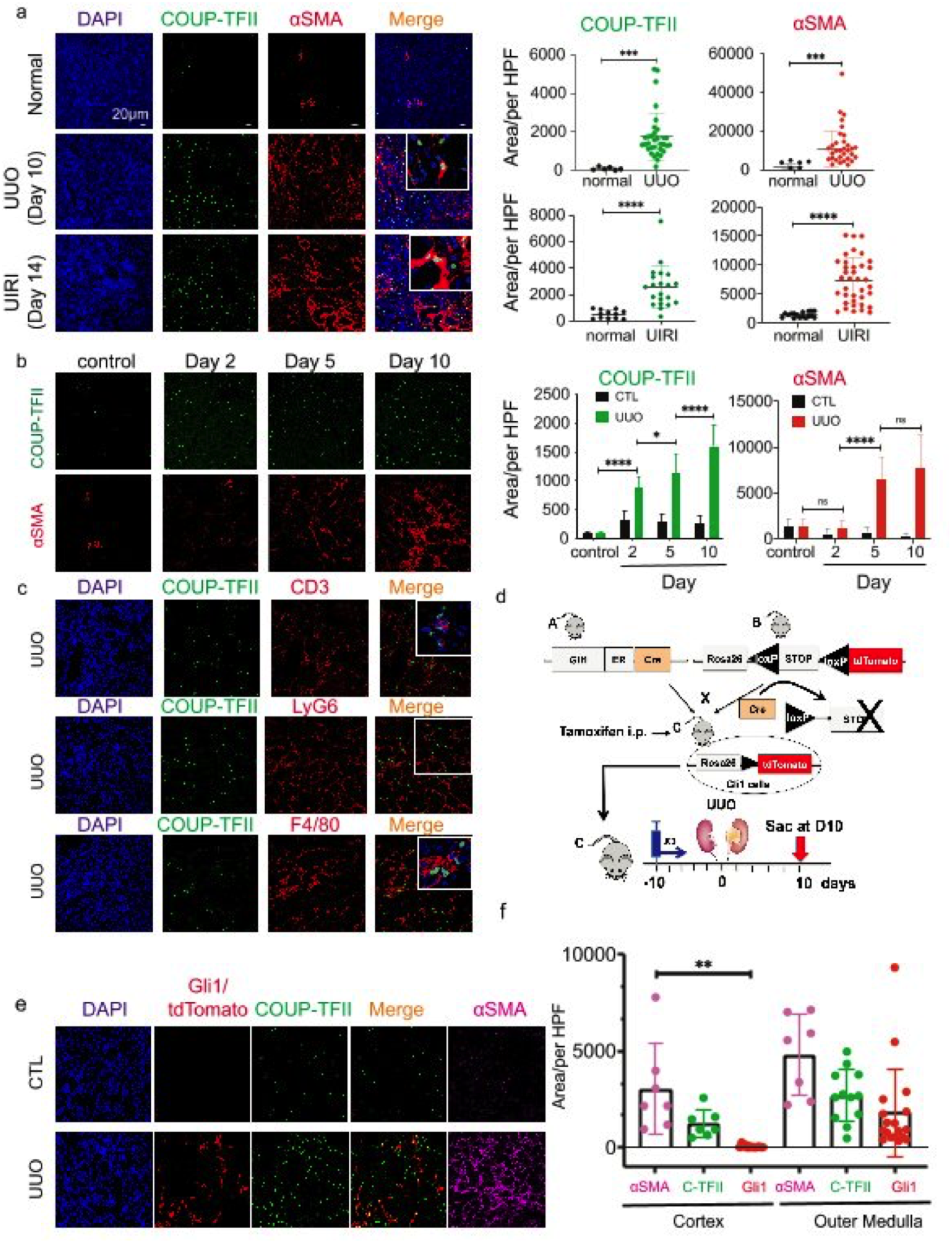
COUP-TFII expression is increased and co-localized with αSMA+ cells within fibrotic regions in the injured mouse kidneys. (a) COUP-TFII expression is significantly increased and co-localized with αSMA+ cells at day 10 of UUO and day 14 of UIRI in mice (n=3 animals per group). Quantification by confocal micrographs in 200x hpf. ***p<0.001 by t test; mean ± SD. (b) In a time course of UUO, COUP-TFII expression increased as early as day 2 and preceded the increased expression of αSMA (n=3 animals per time point). Quantification by confocal micrographs in 200x hpf. ***p<0.001 by t test; mean ± SD. (c) In the UUO model, COUP-TFII positive cells do not stain with markers of inflammatory cells (T cell marker CD3, neutrophil marker LyG6, and macrophage marker F4/80) (n=3 animals). (d) Using a fate tracing of Gli1+ cells in the UUO model, (e&f) *Gli1*-tdTomato+ cells expanded primarily in the outer medullary region. In contrast, COUP-TFII+ cells are distributed both in cortex and medulla. A subset of COUP-TFII+ cells (green) overlap with genetically labeled Gli1+ pericyte/perivascular cells (red) in UUO injury model (n=3 animals per group). CTL: contralateral kidney. Quantification by confocal micrographs in 200x hpf. ***p<0.001 by t test; mean ± SD.

Genetic lineage tracing analysis demonstrated that Gli1 marks perivascular mesenchymal stem cells-like cells, which are proposed to contribute to organ fibrosis (Kramann et al., 2015). We evaluated whether COUP-TFII+ cells and Gli1+ cells represent the same or closely related populations. We used the *Gli1*-CreERt2 line crossed with tdTomato reporter mice and induced genetic labeling 10 days prior to UUO injury (Fig 3d). Kidney tissues were collected at day 10 after UUO and stained for COUP-TFII expression. As expected, *Gli1*-derived cells expanded in number and acquired αSMA expressing 10 days after UUO (Fig 3e). Co-staining revealed that COUP-TFII expression was indeed found in nearly all of Gli1-tdTomato+ cells; however, there were also a large number of COUP-TFII+ cells that were distinct from the Gli1 lineage (Fig 3e). Although some of these cells could result from incomplete labeling with the inducible *Gli1*-Cre, there was also a noticeable difference in the spatial distribution between the COUP-TFII+/tdTomato- and COUP-TFII+/tdTomato+ populations. Interestingly, while COUP-TFII+ cells were found throughout both the cortex and medulla, co-localizing with αSMA+ cells in both regions, the Gli1-tdTomato+ cells were largely restricted to the outer medullary region (Fig 3f). This is consistent with the previous observations (Kramann et al., 2015), that *Gli1*-tdTomato+ cells represented only a small fraction of the total PDGFRβ+ population. Gli1 cells were enriched in the outer medulla with much less expression in pericytes and perivascular fibroblasts of the cortex(Humphreys, 2018, Kramann et al., 2015). As fibrosis is present both in cortex and outer medulla, our data suggest that COUP-TFII functions more generally in regulating pericyte to myofibroblast differentiation, irrespective of the anatomic compartments.

### Ablation of COUP-TFII in adult mice attenuates injury-induced kidney fibrosis

Since global knockout of COUP-TFII in mice confers embryonic arrest at E10 due to defective angiogenesis and cardiac development (Pereira et al., 1999), we used a tamoxifen-inducible Cre-loxP system to knockout COUP-TFII in adult mice. We generated COUP-TFII^flox/+^; Rosa26^CreERT2/+^ (*F*/+; *Cre/+*) and COUP-TFII^flox/flox^; Rosa26^CreERT2/+^ (*F/F*; *Cre/+*) mice. COUP-TFII expression was maintained in these mice without tamoxifen (TAM). Three injections of TAM activated *Cre* recombinase and generated COUP-TFII heterozygous (+/-) and homozygous (-/-) knockout mice. Ten days after TAM injection, mice were subjected to UUO and euthanized 7 days after UUO (Fig 4a). There was no identifiable phenotypic difference between WT and knockout mice without injury.

**Figure 4.**
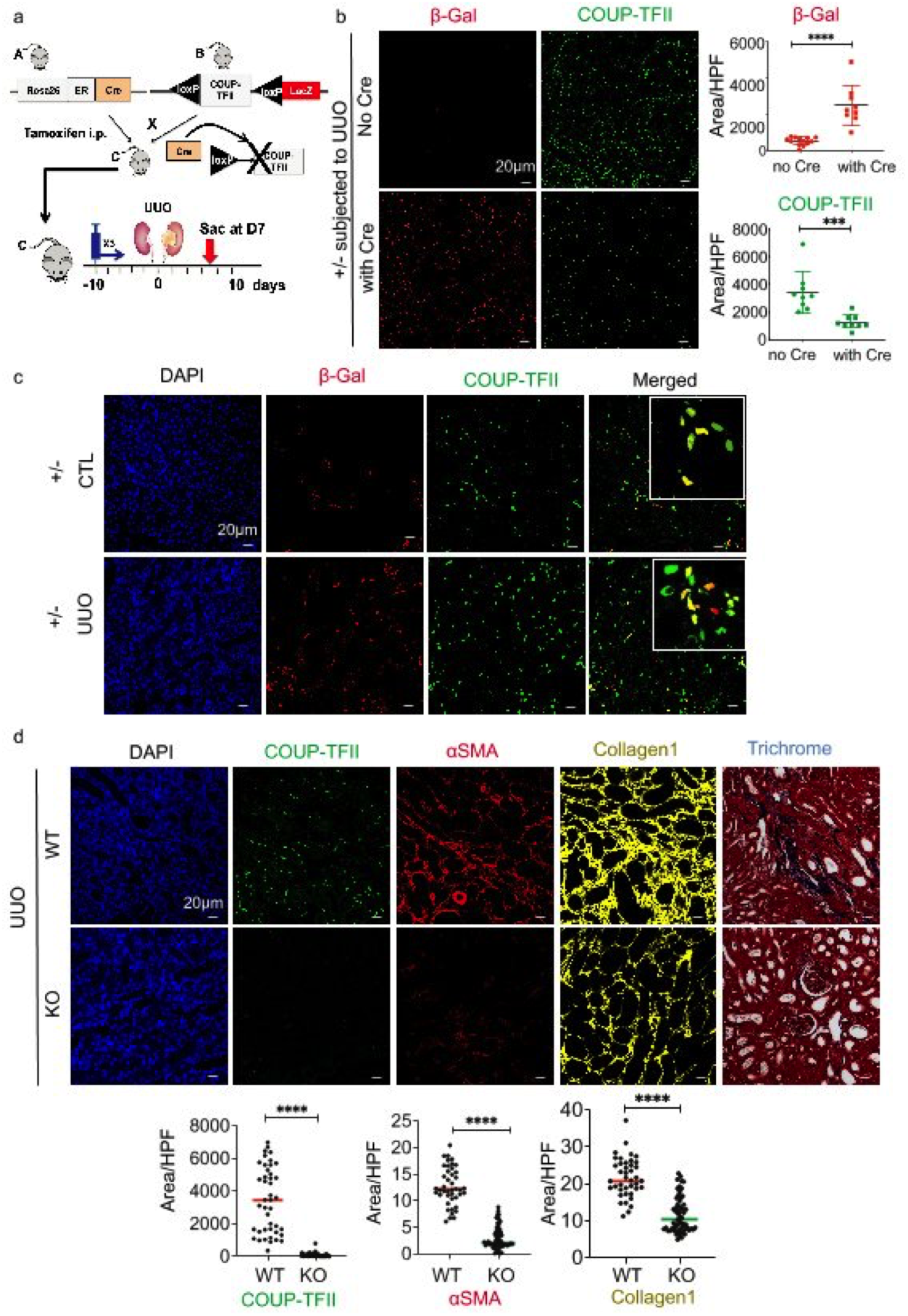
Genetic ablation of COUP-TFII in adult mice attenuates injury-induced kidney fibrosis. (a) Strategy of conditional knockdown of COUP-TFII in adult mice. A *LacZ* knock-in allele is inserted into the genomic COUP-TFII locus after the second *LoxP* site. (b) Activation of Cre recombinase by Tamoxifen (TAM) results in COUP-TFII deletion and the expression of LacZ reporter (detected by immunostaining of β-galactosidase (β-Gal) in the UUO mice model (n=3). Quantification by confocal micrographs in 200x hpf. ***p<0.001 by t test; mean ± SD. (c) Using heterozygous mice (F/+; Cre/+), β-Gal+ cells (red) increased after TAM injection in UUO kidneys, and co-localize with COUP-TFII+ cells (green) (n=3). (d) COUP-TFII+ cells decrease significantly in the UUO kidney in TAM-treated homozygous (F/F;Cre/+) mice (KO group, n=6) compared to wild-type littermates (WT group, n=4). Expression of αSMA (red) and collagen1 (yellow) are also markedly reduced. Masson Trichrome staining shows less kidney fibrosis in KO compared to WT group. Quantification by confocal micrographs in 200x hpf. There are 8-10 images taken and quantified for each animal (represented by each dot). ***p<0.001 by t test, mean ± SD.

We first examined the efficacy of COUP-TFII ablation. A *LacZ* knock-in allele was inserted into the genomic COUP-TFII locus after the second *LoxP* site (Takamoto, You et al., 2005). Treatment with Cre recombinase resulted in COUP-TFII deletion and the expression of the LacZ reporter. Using this *LacZ* reporter, we were able to trace the COUP-TFII lineage following deletion. As shown in Fig 4b, expression of β-galactosidase (β–Gal) increased after TAM injection in UUO kidneys of heterozygous mice with Cre allele, but not in the control mice with flox allele but without Cre allele (WT), indicating the specificity of β-Gal stainin*g* in the *Cre-LoxP* system. Expression of COUP-TFII decreased after TAM injection in heterozygous (+/-) mice. Ten days after TAM injection, β-Gal expression co-localized with COUP-TFII expression in both contralateral and UUO kidneys, consistent with the deletion of one allele of COUP-TFII (Fig 4c). Both β-Gal expression and COUP-TFII expression increased in UUO kidney compared to contralateral non-injured kidney (Fig 4c). β-Gal staining demonstrated that this population remained stable following TAM injection, indicating that COUP-TFII deletion does not impact the survival or proliferation of these cells at baseline.

Next we examined the effect of COUP-TFII ablation on injury-induced kidney fibrosis (UUO model) using knockout (KO, -/-) mice. As shown in Fig 4d, COUP-TFII positive cells decreased significantly in a KO (-/-) mice after UUO compared with WT kidneys, indicating successful knockout of COUP-TFII. Associated with much less COUP-TFII expression, αSMA and collagen1 expression was significantly decreased in KO compared to WT at 7 days after UUO. Histological evaluation demonstrated less kidney fibrosis in KO compared to WT by Masson Trichrome (Fig 4d). These data demonstrated that ablation of COUP-TFII in adult mice attenuates injury-induced kidney fibrosis.

### COUP-TFII regulates myofibroblast differentiation in vitro through metabolic reprogramming

In order to dissect the mechanistic role of COUP-TFII in myofibroblast differentiation, we generated COUP-TFII loss- and gain-of-function cell lines using CRISPR-Cas9 (clustered regularly interspaced short palindromic repeats (CRISPR)–CRISPR associated protein 9 (Cas9)) and an inducible lentiviral construct, respectively, in the pericyte-like cell line C3H/10T1/2. C3H/10T1/2 is a mouse mesenchymal cell that has been used in studies related to pericyte biology and cell fate determination in vitro [45-47]. Naïve C3H/10T1/2 cells (WT) showed basal expression of COUP-TFII, which we were able to successfully modulate using the knockout (KO) or overexpression (OE) systems (Fig 5a). Neither KO nor OE affected cell viability (Fig 5b), although a decreased proliferation rate in KO cells and increased proliferation rate in OE cells were observed compared to naïve (WT) cells (Fig 5c). Treatment of C3H/10T1/2 cells with TGFβ1 induces myofibroblast differentiation with up-regulation of αSMA, and this effect was preserved in KO and OE cells (Fig 5d). Interestingly, OE cells, in the absence of TGFβ1 stimulation, displayed an elongated fibroblast shape, similar to WT cells treated with TGFβ1 (Fig 5d). More importantly, there was increased expression of αSMA and collagen1 in COUP-TFII OE cells even in the absence of TGFβ1 stimulation (Fig 5d-e), while their expression was significantly diminished in the KO cells. Collectively, these data further support a profibrotic function for COUP-TFII in pericyte/fibroblast-like cells.

**Figure 5.**
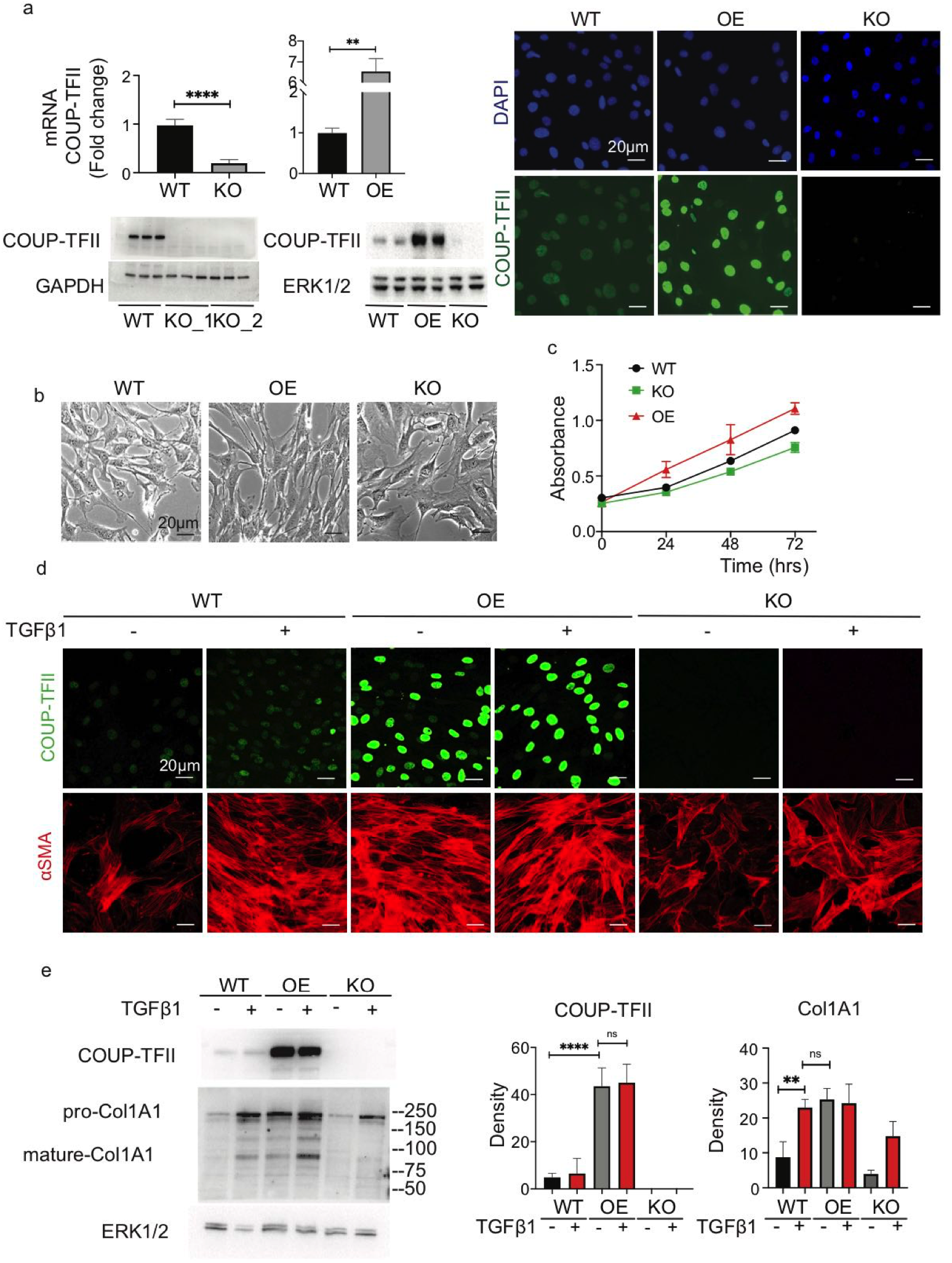
Cellular knockout of COUP-TFII *in vitro* in C3H/10T1/2 cells decreases cell proliferation and suppresses TGFβ1-induced αSMA and collagen1 expression. In contrast, overexpression of COUP-TFII alone increases collagen1 production. (a) Verification of COUP-TFII loss- and gain-of-function cell lines generated by CRISPR-Cas9 and an inducible lentiviral construct in pericyte-like cell line C3H/10T1/2 *in vitro* (n=6). (b&c) COUP-TFII KO had no effect on cell viability, although decreased proliferation rate compared to naïve C3H/10T1/2 cells (WT) (n=6), ****p<0.0001 by two-way ANOVA. (d) COUP-TFII-OE cells, in the absence of TGFβ1 stimulation, displayed an elongated fibroblast shape, similar to WT cells treated with TGFβ1. (e) Overexpression of COUP-TFII alone without TGFβ1 induces collagen1 production (n=3). ****p<0.0001 by one-way ANOVA, mean ± SD.

To further interrogate the molecular mechanisms by which COUP-TFII regulates myofibroblast differentiation, we evaluated the proteome of WT and OE cells that were differentiated with TGFβ1. This non-biased approach unequivocally implicated cellular metabolic pathways as predominant targets of COUP-TFII. Gene set enrichment analysis (GSEA) highlighted that the most significantly down-regulated proteins in COUP-TFII-OE cells were enriched in mitochondrial electron transport chain and fatty acid oxidation (FAO) pathways (Fig 6a), which are involved in oxidative metabolism. Conversely, the top up-regulated pathways in COUP-TFII-OE cells were associated with both extracellular matrix, including collagen fiber organization, cadherin binding and actin cytoskeleton, and glycolysis pathways (Fig 6a & b). Taken together, these data support a model in which COUP-TFII shifts cell metabolism from FAO toward glycolysis, which may lead to increased extracellular matrix production and fibrosis. Indeed we found that COUP-TFII overexpression was sufficient to significantly increase expression of key components of the glycolytic pathway, including Glut1, Hexokinase2 (HK2) and Lactate dehydrogenase A (LDHA) (Fig 6c). To directly test the link between glycolytic metabolism and myofibroblast differentiation, we inhibited glycolysis with 2-deoxyglucose (2-DG), a well-defined HK2 inhibitor. In the COUP-TFII OE cells, 2-DG attenuated the TGFβ1-induced expression of αSMA and collagen1 at both the mRNA and protein level (Fig 6d). These data suggest that COUP-TFII enhances TGFβ1-induced glycolysis during myofibroblast differentiation, and that the switch to glycolysis is essential for this process.

**Figure 6.**
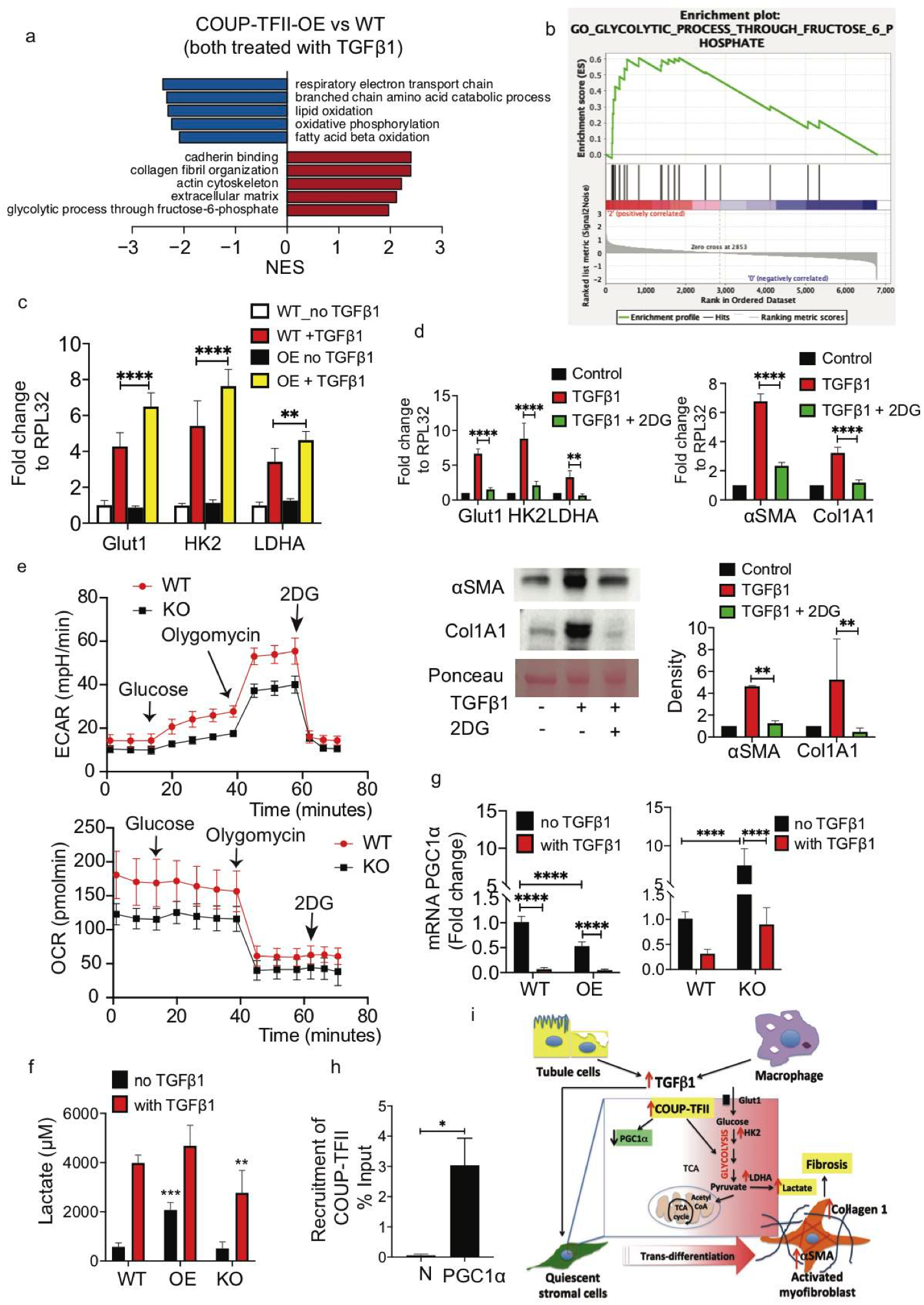
COUP-TFII regulates myofibroblast differentiation through metabolic reprogramming *in vitro.* (a&b) Non-biased proteomics of TGFβ1-treated naïve (WT) and COUP-TFII overexpressing (COUP-TFII-OE) cells reveals a metabolic shift from fatty acid oxidation to glycolysis. Top pathways ranked by NES (gene set enrichment with FDR < 0.05) altered in COUP-TFII-OE compared to naïve C3H10T1/2 cells (n=2). (c) qRT–PCR of glycolysis genes for WT and COUP-TFII-OE cells treated without or with TGFβ1 (10ng/ml) (n=6). **p<0.01, ****p<0.0001 by two-way ANOVA. (d) 2-deoxyglucose (2-DG) treatment inhibits TGFβ1-induced glycolysis (left) and fibrosis (right) in COUP-TFII-OE cells by qRT-PCR (n=6). Collagen1 protein is reduced by 2-DG confirmed by western blot (n=3). ****p<0.0001 by oneway ANOVA. (e) WT and COUP-TFII knockout (COUP-TFII-KO) cells were seeded in SeahorseXF-24 cell culture microplates. The cells were rendered quiescent in 1% BSA DMEM overnight and then treated with 10 ng/ml TGFβ1 for 24 hours, followed by sequential treatments with glucose (10mM), oligomycin (1 μg/ml) and 2-DG (100 mM). Real-time extracellular acidification rate (ECAR) was recorded (n = 10). When compared with WT, COUP-TFII-KO cells significantly decreased ECAR induced by TGFβ1. Similar difference were seen after oligomycin injection, which indicated the glycolytic capacity. ****p<0.0001 by t test. Interestingly, KO cells displayed a lower oxygen consumption rate (OCR), a measure of mitochondrial respiratory activity, in response to TGFβ1 compared with WT cells. (f) TGFβ1 significantly increased more lactate production by WT C3H/10T1/2 cells when compared to COUP-TFII-KO cells treated with TGFβ1. In contrast, overexpression of COUP-TFII alone (without TGFβ1) significantly increased lactate (n=3). **p<0.01, ***p<0.001 by one-way ANOVA, mean ± SD. (g) Overexpression of COUP-TFII inhibited, and knockout of COUP-TFII increased, PGC1α mRNA (n=6). ***p<0.001 by one-way ANOVA, mean ± SD. (h) Chip-qPCR analysis on C3H10T1/2 cells *in vitro* revealed binding of COUP-TFII on the promoter of *PGC1α* (n=2). *p <0.05, mean ± SD. (i) Schematic model of the role of COUP-TFII in metabolic reprogramming during myofibroblast differentiation and fibrosis formation after injury.

Having demonstrated that COUP-TFII-OE cells adopt enhanced glycolysis and profibrotic phenotype in response to TGFβ1, we further examined this model in the KO cell line. Using the Seahorse X24 extracellular flux analyzer, we found that the COUP-TFII KO cells exhibited significant reduction in both the glucose-induced glycolytic preference and the oligomycin-induced total glycolytic capacity when compared to WT cells, measured by extracellular acidification rate (ECAR) (Fig 6e). Interestingly, KO cells has a lower oxygen consumption rate (OCR), a measure of mitochondrial respiratory activity, in response to TGFβ1 compared with WT cells (Fig 6e). This is consistent with the slower growth of KO cells compared to WT cells. To confirm modifications in glycolysis induced by COUP-TFII perturbation, we directly measured the levels of lactate in the culture medium. TGFβ1 significantly increased lactate production in WT cells, which was blunted by COUP-TFII KO. Overexpression of COUP-TFII alone (without TGFβ1) significantly increased lactate levels (Fig 6f), indicating augmented glycolysis in these cells. Furthermore, overexpressing COUP-TFII resulted in decreased transcription of PGC1α and knockout of COUP-TFII increased PGC1α mRNA in vitro (Fig 6g). This is consistent with our Chip-qPCR results, in which PGC1α is a target of COUP-TFII (Fig 6h), Altogether, these data demonstrated that COUP-TFII augmented glycolysis and promoted myofibroblast differentiation, and collagen1 production after injury (Fig 6i).

## Discussion

Pericyte/perivascular cells are important contributors to kidney fibrosis after injury (Humphreys et al., 2010, Lin et al., 2008). These cells are the main source of myofibroblasts, the effector cells for fibrosis (Duffield, 2014, Humphreys, 2018). Blocking the differentiation of pericyte/perivascular cells to myofibroblasts is an attractive strategy to reduce fibrosis. TGFβ, a master regulator of myofibroblast differentiation, has been extensively studied as a therapeutic target (Akhurst & Hata, 2012, Meng, Huang et al., 2012, Rangarajan, Kurundkar et al., 2016). Direct inhibition of TGFβ, however, has led to more toxicity than benefit (Li, Wan et al., 2006, Principe, Doll et al., 2014). Therefore, alternative molecular approaches to the regulation of myofibroblast differentiation during fibrosis development are desirable for anti-fibrotic drug development.

We demonstrated that COUP-TFII is markedly increased in human kidneys and lungs with fibrosis or cells in human kidney organoids activated with IL-1β to enhance stromal fibrosis. COUP-TFII is expressed in pericytes/perivascular cells in adult non-injured kidney, and colocalizes with αSMA expression during fibrosis formation after kidney injury. Another protein proposed to be important in the conversion of pericytes/perivascular cells to myofibroblasts is Gli1 (Kramann et al., 2015). In comparison with Gli1, COUP-TFII is seen in more αSMA+ cells, including many that are not *Gli1*-tdTomato+, especially in the kidney cortex. Increased COUP-TFII expression after injury has important functional significance for the myofibroblast population. Using a tamoxifen-induced *Cre*-loxP system, we deleted COUP-TFII in adult mice and evaluated its role in injury-induced kidney fibrosis. Genetic depletion of COUP-TFII reduced αSMA positive cells and attenuated kidney fibrosis after injury. Our results demonstrate that COUP-TFII plays a pivotal role in myofibroblast differentiation and kidney fibrosis formation. We also demonstrated increased COUP-TFII expression in myofibroblasts in fibrotic lungs of patients with idiopathic lung fibrosis (IPF). These findings implicate the general role of COUP-TFII in myofibroblast during fibrosis formation.

Giving the important role of COUP-TFII on glucose and lipid metabolism (Ashraf et al., 2019), we hypothesized that COUP-TFII might regulate myofibroblast differentiation through metabolism reprogramming. Our proteomic data revealed that COUP-TFII promotes the expression of proteins enriched in metabolic process critical for myofibroblasts, in particular, suppression of FAO and enhancement of glycolysis. Our data link COUP-TFII to the metabolic switch from oxidative metabolism to aerobic glycolysis leading to a proliferative and pro-fibrotic function of myofibroblast. Overexpression of COUP-TFII alone (without TGFβ1 treatment) is sufficient to increase glycolysis and collagen1 expression. Knockout of COUP-TFII dampened TGFβ1-induced glycolysis and decreased αSMA and collagen1 expression. The phenotypic resemblance between COUP-TFII overexpressed alone cells and WT cells treated with TGFβ1 suggests that COUP-TFII is essential to establish the myofibroblast phenotype through augmented glycolysis. High glycolytic flux is important for the self-renewal of progenitor cells (Liu, Edgington-Giordano et al., 2017). Interestingly, COUP-TFII expression is abundant in the mesenchymal compartment of the developing organs during embryonic organogenesis but declines significantly right after birth(Pereira, Qiu et al., 1995). It is tempting to suggest that COUP-TFII might be important to maintain the de-differentiation status of cells through augmented glycolysis during development.

Besides supporting energy needs in a hypoxic environment, glycolysis also provides macromolecules required for cell proliferation and migration (Xie et al., 2015). In this way, the reprogramming of cell metabolism ensures sufficient building blocks for biosynthesis and facilitates survival of myofibroblasts in a harsh hypoxic and nutrient-deprived microenvironment. Furthermore, glycolysis is clearly linked to ECM production (Ding et al., 2017, Zhao et al., 2019). Collagen1, the predominant structural protein found in kidney fibrosis, is synthesized through multiple steps, including hydroxylation of amino acids, disulfide bonding and glycosylation (Basak, Vega-Montoto et al., 2016). The major amino acid components of collagen are glycine, proline, and lysine. Glycine is mainly produced from glycolysis (de Paz-Lugo, Lupianez et al., 2018). In addition, collagen hydroxylation and glycosylation are depended on glycolysis (Im, Freshwater et al., 1976). Our data, along with studies from others (Ding et al., 2017, Xie et al., 2015), demonstrate that 2-DG (an hexokinase inhibitor) drastically decreases collagen1 production in myofibroblasts *in vitro*. There is emerging evidence showing that augmented glycolysis in cancer stromal cells (also call cancer associated fibroblasts) support cancer progression through secreting lactate and other glycolytic intermediates (Avagliano, Granato et al., 2018). In addition, a decrease in microenviromental pH from lactic acid accumulation has been associated with increased TGFβ activity (Kottmann, Kulkarni et al., 2012). Therefore, targeting glycolysis in myofibroblasts would be expected to not only inhibit the activation of stromal cells, but also to modify the microenvironment of fibrosis foci. We report that this molecular regulation of metabolic reprogramming is likely facilitated by the function of COUP-TFII to decrease PGC1α transcription given that COUP-TFII binds to the PGC1α promoter.

Our study shows that with fibrosis COUP-TFII is expressed primarily in myofibroblasts. Since COUP-TFII is expressed primarily in myofibroblasts, targeting it may be selectively effective to suppress myofibroblast differentiation and function without affecting epithelial or endothelial cells. There are no overt phenotypes in COUP-TFII KO mice at the adult stage further supporting the potential safety profile of COUP-TFII inhibition. Given that COUP-TFII expression is upregulated in fibrotic human kidney organoids, these ex vivo human cell systems can be used to test potential inhibitors.

In conclusion, our study provides compelling evidence that COUP-TFII regulates myofibroblast differentiation and profibrotic function through augmented glycolysis after injury. Reducing COUP-TFII is effective in diminishing myofibroblast differentiation and limiting fibrosis. The fibrogenic response may share a common pathway in different organ injury and failure. Targeting COUP-TFII serves as a novel treatment approach for mitigating fibrosis in chronic kidney disease and potential other organ fibrosis.

## Methods and Material

### Human kidney biopsy sample preparation and immunostaining

Normal kidney samples used for immunostaining studies were obtained from surgical sections from patients undergoing nephrectomy due to renal cell carcinoma (RCC) under institutional review board-approved protocols. Injured kidney samples were obtained form kidney biopsy samples from patients with thrombotic microangiopathy (TMA) and diabetic nephropathy (DN).

### Human kidney organoids generation and IL-1β stimulation

Kidney organoids were derived from H9 human embryonic stem cells as previously described (Morizane et al., 2015). Briefly, H9 cells were cultured in StemFit (Ajinomoto) supplemented with 10 ng/ml FGF2 (Peprotech). Differentiation was started with 8 υM CHIR (TOCRIS) for 4 days, followed sequentially by Activin for 3 days and FGF9 for 1-2 days. Subsequently cells were dissociated with Accutase (Stem Cell Technologies) and plated into U-shaped bottoms 96-well plates (Corning) at 100,000 cells per well in a medium supplemented with 3 μM CHIR and 10 ng/ml FGF9. Two days later the medium was changed to one supplemented with only 10 ng/ml FGF9. After 3-4 days, the medium was changed to basal medium without additional growth factors. At day 51, matured organoids were treated with IL-1β 10 ng/ml (Sigma) for 96 hours. Organoids were collected and fixed in 4% PFA for 30 minutes followed by 20% sucrose overnight. Cryosections (7 μm) were incubated with antibodies for immunofluorescence. Images were captured by confocal microscopy using a Nikon C1 microscope running EZ-C1 software.

### Mouse strain and animal experiments

All mouse experiments were performed under the animal use protocol approved by the Institutional Animal Care and Use Committee of the Brigham and Women’s hospital. *Gli1-* CreER^t2^ (JAX# 007913), *Rosa26*tdTomato (JAX# 007909), *Foxd1*-GFP-Cre (Jax# 012463), Rosa26-CreER^t2^ (JAX# 008463) were purchased from Jackson Laboratories (Bar harbor, ME). COUP-TFII flox/+ mice were purchased from Mutant Mouse Resource & Research Centers (MMRRC) (B6;129S7-Nr2f2tm2Tsa/Mmmh, Cat# 032805-MU). The COUP-TFII flox/+ mouse strain was maintained in a mixed genetic background (129/Sv x C57BL/6) and received standard rodent chow. To induce COUP-TFII deletion in the adult, 8-12 weeks old mice were intraperitoneally injected with 3 doses 0.1 mg/g body weight tamoxifen in corn oil/3% ethanol (Sigma) every other day starting 14 days before surgery.

Murine kidney fibrosis models were performed as previously described (Yang, Besschetnova et al., 2010). Briefly, mice were anesthetized with pentobarbital sodium (60 mg/kg body weight, intraperitoneally). For the unilateral ureteral obstruction (UUO) surgery, a flank incision was made and the left ureter was tied off at the level of the lower pole with two 4.0 silk ties. For the unilateral ischemia reperfusion injury (IRI), the left kidney was exposed through a flank incision and subjected to ischemia by clamping the renal pedicle with non-traumatic microaneurysm clamps (Roboz, Rockville, MD) for 30 minutes. Reperfusion was confirmed by visual color change. Body temperatures were controlled at 36.5°C–37.5°C throughout the procedure. One milliliter of warm (body temperature) saline was instilled in the retroperitoneum after surgery for volume supplement. Buprenorphine was used for pain control (0.1mg/kg body weight, intraperitoneally). Mice were sacrificed at day 2, 5 or 10 after UUO and day 14 after unilateral IRI.

### Histology and immunofluorescence staining

Mice were anesthetized with isofluorane (Baxter) and subsequently perfused via the left ventricle with 4°C PBS for 1 minute. Kidneys from adult mice were fixed with 4% paraformaldehyde (PFA), dehydrated and embedded in paraffin. Hematoxylin/eosin, PAS and Masson’s trichrome staining were performed using standard protocols (Yang et al., 2010).

Immunofluorescence staining of mouse kidneys was performed on paraffin sections as previously described (Yang et al., 2010). Briefly, the tissue sections were deparaffinized, followed by antigen retrieval and rehydration. Then tissue antigens were labeled with primary antibodies to COUP-TFII (Abcam, diluted 1: 200), αSMA (Sigma, 1:400), collagen1 (EMD Millipore, 1:400), CD31 (Abcam 1: 200), PDGF receptor beta (PDGFRβ, Abcam 1:400), or β-galactosidase (Abcam 1:100), followed by FITC or Cy3-labeled secondary antibodies (Jackson ImmunoResearch). Some immunostaining was performed on frozen sections. Cryosections (7 μm) were fixed in 4% PFA for 2 hours, and then, washed in 30% sucrose solution overnight. Primary antibodies used in cryosections recognized the following proteins: CD3 (eBioscience 1:100), LyG6, F4/80 (ThermoFisher), Images were captured by a confocal (Nikon C1) or standard fluorescent microscope (Nikon TE 1000).

### Cell Culture and treatment

The C3H10T1/2 (American Type Culture Collection) were cultured in DMEM medium supplemented with 10% FCS until the cells were 80% confluent. Cell morphology was examined and captured using live cell imaging (Nikon). For cytokine induction of myofibroblast differentiation, subconfluent cells were incubated in DMEM medium containing 1% BSA overnight and treated with TGFβ1 (10ng/ml) (R&D).

### CRISPR/Cas9 knockout

COUP-TFII guide RNAs were created using a guide design web tool (http://crispr.mit.edu): COUP-TFII guide 1, TATATCCGGACAGGTACGAG; COUP-TFII guide 2, GAGGGGGTCCCCGTTGGTCA. sgRNA oligos were cloned into pSpCas9(BB)-2A-GFP (Addgene, 48138). The protocol was performed according to published methods (Ran, Hsu et al., 2013). C3H10T1/2 cells (5 x 10^4^) were transfected with lipofectamine 3000. Cells were grown for 48 h and then sorted for GFP and positive cells seeded as single cells in 96-well plates. Cells were then returned to the incubator and cultures were allowed to expand for 2-3 weeks.

### Tet-Inducible COUP-TFII expression

To generate a lentiviral transfer plasmid for inducible gain-of-function experiments, we used high-fidelity PCR (iProof, BioRad) to amplify full-length mouse COUP-TFII cDNA, flanked with attB1 and attB2 sites on 5’ and 3’ ends respectively, from a cDNA library from adult mouse kidney. The resulting PCR band was purified from a 0.8% agarose gel using QIAquick Gel Extraction kit (Qiagen). The purified PCR product was then cloned into pDONR221 (Thermo Fisher Scientific) using BP Clonase II (Invitrogen) according to manufacturer’s instructions. We then shuttled the COUP-TFII cDNA into the destination vector pInducer20 (a gift from Stephen Elledge, Addgene #44010) using LR Clonase II (Invitrogen). All cloning steps were verified using Sanger sequencing (Genewiz, Inc.). C3H10T1/2 cells were infected with lentivirus in the presence of 10 μg/ml polybrene (Sigma). Infected cells were selected with puromycin (Sigma).

### Cell proliferation assay

Cells were seeded on a 96-well plate at a concentration of 1 × 10^4^ per well. Three parallel wells of cells were studied for each group. After incubation for 1 day, 2 days, or 3 days, 20 μL MTT ((3-(4,5-Dimethyl-2-thiazolyl)-2,5-diphenyltetrazolium Bromide), promega) was added. After 2 hours of MTT exposure, cells were washed and subjected for colorimetric measurement. The OD values were obtained by a microplate reader (SpectraMax M5, Molecular Devices) at 570 nm.

### Quantitative real-time PCR (qRT-PCR)

At indicated times, total RNA was extracted using TRIzol (Sigma) as described (Yang et al., 2010). Subsequently, 2 μg of total RNAs was reverse-transcribed to cDNA with random primers using reverse transcriptase (Invitrogen). A 1: 5 dilution of cDNA was then amplified by real-time qPCR in a CFX96 real-time system (Biorad) using SYBR green. Relative gene expression was calculated by the ΔΔCt method, and final results were expressed as the fold difference relative to control conditions in gene expression normalized to Ribosomal Protein L32 (RPL32). Primers for individual gene expression were listed in Table 1.

**Table 1.**
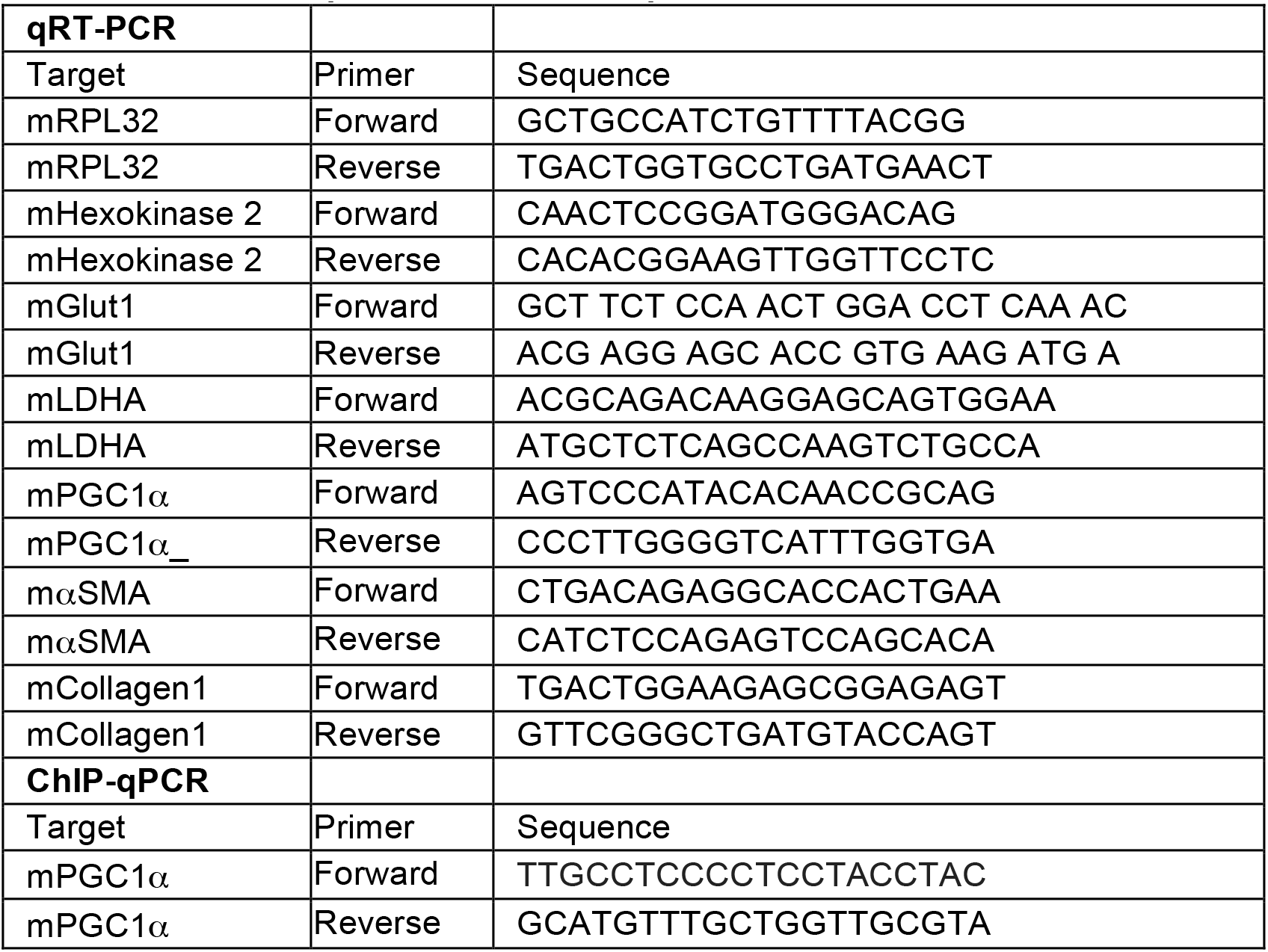
Primers for, qRT-PCR and ChIP-qPCR.

### ChIP assay

Chromatin immunoprecipitation and real-time PCR quantification were performed as described (Mukhopadhyay, Deplancke et al., 2008). The rabbit polyclonal COUP-TFII antibody (Millipore, Cat# ABE 1426) and rabbit IgG1 antibody (Zymed) were used for immunoprecipitation. After purification of DNA from C3H10T1/2 cells, bound sequences were determined by quantitative real-time PCR (Table 1 for the primer sequences for the ChIP assay).

### Western Blot Analysis

48 hours after transfection, C3H10T1/2 cells were lysed with 1xRIPA buffer containing a protease inhibitor cocktail (Roche Applied Science) for 30 minutes on ice. After centrifugation for 15 minutes, the supernatant was collected, and protein content of the samples was analyzed according to the Bradford method. Proteins were loaded onto SDS-polyacrylamide gels and blotted onto PVDF membrane (Bio-Rad Laboratories). Western blots were performed using antibodies directed against COUP-TFII (abcam, 1: 2000), αSMA (Sigma, 1:4000), and ERK2 (Cell signaling). HRP-conjugated secondary antibodies were purchased from DAKO. Enhanced chemiluminescence was performed according to the manufacturer’s instructions. ERK1/2 protein was used to ensure equivalent loading of protein samples.

### Proteomics

Naïve (WT) and COUP-TFII overexpression (OE) cells were both treated with TGFβ1 (10ng/ml) for 48 hours to induce myofibroblast differentiation. After trypsinization, cells were washed twice in PBS. Cell pellets were frozen in – 80 C° for proteomics experiments.

#### Protein extraction and digest

PBS was removed, followed by addition of SDS lysis buffer (2% SDS, 150 mM NaCl, 50 mM Tris, pH 8.7) containing protease inhibitors (Complete, Roche). Lysates were homogenized over Qiashredder columns (Qiagen, ref. 79656) and centrifuged at 13,000 rpm for 1 minute at room temperature. Reductive methylation of cysteine residues was performed by adding dithiothreitol (DTT) to a final concentration of 5 mM and heating to 37 C° for 1 hour, followed by alkylation with iodoacetamide at a final concentration of 15 mM and incubation at room temperature in the dark for 30 minutes and quenching with DTT. Protein concentration was determined using a Micro BCATM Protein Assay Kit (ThermoFisher, Catalog# 23235). Detergent was removed by methanol/chloroform protein precipitation as described previously (Wessel & Flugge, 1984). Lys-C protease digests (Wako, Catalog# 129-02541) in 2M urea 20mM EPPS, pH 8.5 at 37°C for 3 hours were followed by further digestion at 37 °C for 6 hours with trypsin (Promega, Catalog# V5113). Missed cleavage rate was determined by LC-MS/MS.

#### Tandem Mass Tag (TMT) Labeling, Ratio Check and HPLC Fractionation

Equal amounts of protein were removed from each sample and labeled using a TMT11plex Mass Tag Labeling Kit (ThermoFisher, Catalog# A34808). TMT labeling efficiency and ratio checks were determined by LC-MS3 analysis. Quenched TMT labeling reactions were combined and de-salted using a SepPak tC18 Vac RC Cartridge (50 mg, Waters, Catalog# WAT054960). HPLC fractionation was performed using an Agilent 1200 Series instrument with a flow rate of 600 μl/minute over a period of 75 minutes. Peptides were collected in a 96-well plate over a 65 min-gradient of 13-44 %B with Buffer A comprising 5% acetonitrile, 10 mM ammonium bicarbonate, pH 8 and Buffer B comprising 90% acetonitrile, 10 mM ammonium bicarbonate, pH 8. Fractions were then pooled into 24 samples, followed by sample cleanup using the Stage Tip protocol. This protocol uses C18 EmporeTM Extraction Disks (Fisher Scientific, Catalog# 14-386-2). Samples were dried before re-suspension in MS Loading Buffer (3% acetonitrile, 5% FA).

#### LC-MS

Peptides were separated over a 30 cm, 100 μm (internal diameter) column using an EASY-nLC 1200 HPLC system. Samples from the HPLC were injected into an Orbitrap Fusion Lumos Tribrid MS (ThermoFisher, Catalog# FSN02-10000) and measured using a multinotch MS3 method (McAlister, Nusinow et al., 2014, Ting, Rad et al., 2011). MS scans were performed in the Orbitrap over a scan range of 400-1400 m/z. The top 10 ions with charge states from 2 to 6 were selected for MS/MS. Turbo rate scans were performed in the Ion Trap with a collision energy of 35% and a maximum injection time of 250 ms. TMT quantification was performed using SPS-MS3 in the Orbitrap with a scan range of 100-1000 m/z and an HCD collision energy of 55%. Orbitrap resolution was 50,000 (dimensionless units) with a maximum injection time of 300 ms.

#### Proteomic data analysis

Raw data were converted to mzXML format and peptide ID used Sequest (Eng, McCormack et al., 1994) (version 28 (http://fields.scripps.edu/yates/wp/?page_id=17)) with searches against the Human UniProt database (February 2014). The database search included reversed protein sequences and known contaminants such as human keratins that were excluded for subsequent analyses. Linear discriminant analysis was performed (Elias & Gygi, 2007) and peptide false discovery rate (FDR) was < 1% after applying a target-decoy database search strategy. Filtering was performed as described previously (McAlister et al., 2014). Variable modification for oxidized methionine (+15.99 Da) was used during searches. For protein identification and quantification, shared peptides were collapsed into the minimally sufficient number of proteins using rules of parsimony. Peptides with a total TMT value of > 200 and an isolation specificity of > 0.7 were included for quantification.

#### Gene Set Enrichment Analysis (GSEA)

We analyzed our proteome datasets from four independent experiments (WT and COUP-TFII-OE, all in duplicates) using the GSEA software developed by Broad Institute (Cambridge, MA) (Subramanian, Tamayo et al., 2005). All the analyses were performed in default setting based on all GO terms (c5.all.v7.0.symbols.gmt).

### Real-time cell metabolism assay

Naïve (WT) and COUP-TFII KO C3H/10T1/2 cells were plated in XF-24 Cell Culture Microplates (Seahorse Bioscience) at a cellular density of 20,000 cells per well, then serum starved for 24 hours and stimulated with or without TGFβ1 (10 ng/ml) for 24 hours. Real-time analysis of extracellular acidification rate (ECAR) was analyzed using XF Extracellular Flux Analyzer (Seahorse Bioscience). The cells were incubated in basal media followed by sequential injections with glucose (10mM), oligomycin (1 μg/ml) and 2-DG (100 mM).

### Extracellular Lactate Assays

Extracellular levels of lactate were determined using the lactate assay kit (BioVision, Milpitas, CA) according to the manufacturer’s instructions.

### Statistical analysis

Results are expressed as mean ± SD for three experiments done in triplicate or quadruplicate. Group means were compared by one-way analysis of variance (ANOVA), followed by the Tukey post-test using GraphPad Prism (GraphPad software) for multiple comparisons or by student’s t-test. A *p* value <0.05 was considered statistically significant.

## Acknowledgments

This work was supported by the National Institutes of Health Grants R37DK039773, R01DKD072381 and UH3TR002155 (to J.V.B.), National Institute of Biomedical Imaging and Bioengineering (NIBIB), Organ Design and Engineering Training Grant 1T32EB016652-01A1 (to L.L). P.G received support from Monahan Foundation, Fondation pour la Recherche Médicale, Groupe Pasteur Mutualité, Société Francophone de Transplantation, Arthur Sachs fellowship, Philippe Foundation, Fulbright Scholarship, ATIP Avenir program. X.X was supported by China Scholarship Council fellowship. We thank Dr. Edy Kim from Brigham women’s hospital provided human idiopathic lung disease samples for immunostaining.

## Author contributors

LL, PG, XX, ACF, DT, MK and XS performed experiments. LL and JVB designed and analyzed experiments. LL wrote the manuscript.

